# Techniques for visualizing fibroblast-vessel interactions in the developing and adult CNS

**DOI:** 10.1101/2021.12.28.474304

**Authors:** Hannah E. Jones, Kelsey A. Abrams, Julie Siegenthaler

## Abstract

Fibroblasts are found associated with blood vessels in various locations across the CNS: in the meninges, the choroid plexus, and in the parenchyma within perivascular spaces. CNS fibroblasts have been characterized using transcriptional profiling and a *Col1a1-GFP* mouse line used to identify CNS fibroblasts *in vivo*. However, current methods for visualizing CNS fibroblasts are lacking and, in particular, prevent adequate assessment of fibroblast-vessel interactions. Here, we describe methods for whole mount visualization of meningeal and choroid plexus fibroblasts, and optical tissue clearing methods for visualization of parenchymal vessel-associated fibroblasts. Importantly, these techniques can be combined with immunohistochemistry methods for labeling different cell types in the meninges and blood vasculature as well as EdU-based cell proliferation assays. These methods are ideal for visualization of vessel-fibroblast interactions in these CNS structures and provide significant improvement over traditional sectioning and staining methods. We expect these methods will advance studies of CNS fibroblast development and functions in homeostasis, injury, and disease.

## 1. Introduction

The central nervous system (CNS) is comprised of a diverse array of different cell types each with specialized functions. Neuronal, glial, immune, and vascular cell types and sub-types have all been well-profiled, and the mechanisms controlling their development and functions have been robustly studied. More recently, owing in part to single-cell RNA-seqencing studies of CNS cell types, a little-studied population of cells in the CNS has been more thoroughly characterized in mice and humans: fibroblasts. Fibroblasts are a major source of extracellular matrix proteins in tissues and organs^1^ and can have signaling functions (ex: cytokine or growth factor secretion) in development and in health^2^, as well as following injury or in disease^3–5^. In the CNS, fibroblasts are located within close proximity to blood vessels in the meninges, the stroma of the choroid plexus, and in perivascular spaces within the brain parenchyma, and are fluorescently labeled in the *Collagen1a1-GFP* (*Col1a1-GFP*) transgenic mouse line (Figure 1)^6–8^. Not much is known about CNS fibroblast origins during development, heterogeneity, functions, or how they interact with other CNS cell types, namely vascular cell types. Considering the important homeostatic and disease roles of fibroblasts in non-CNS tissues and organs, this makes CNS fibroblasts of particular importance to study.

**Figure 1:**
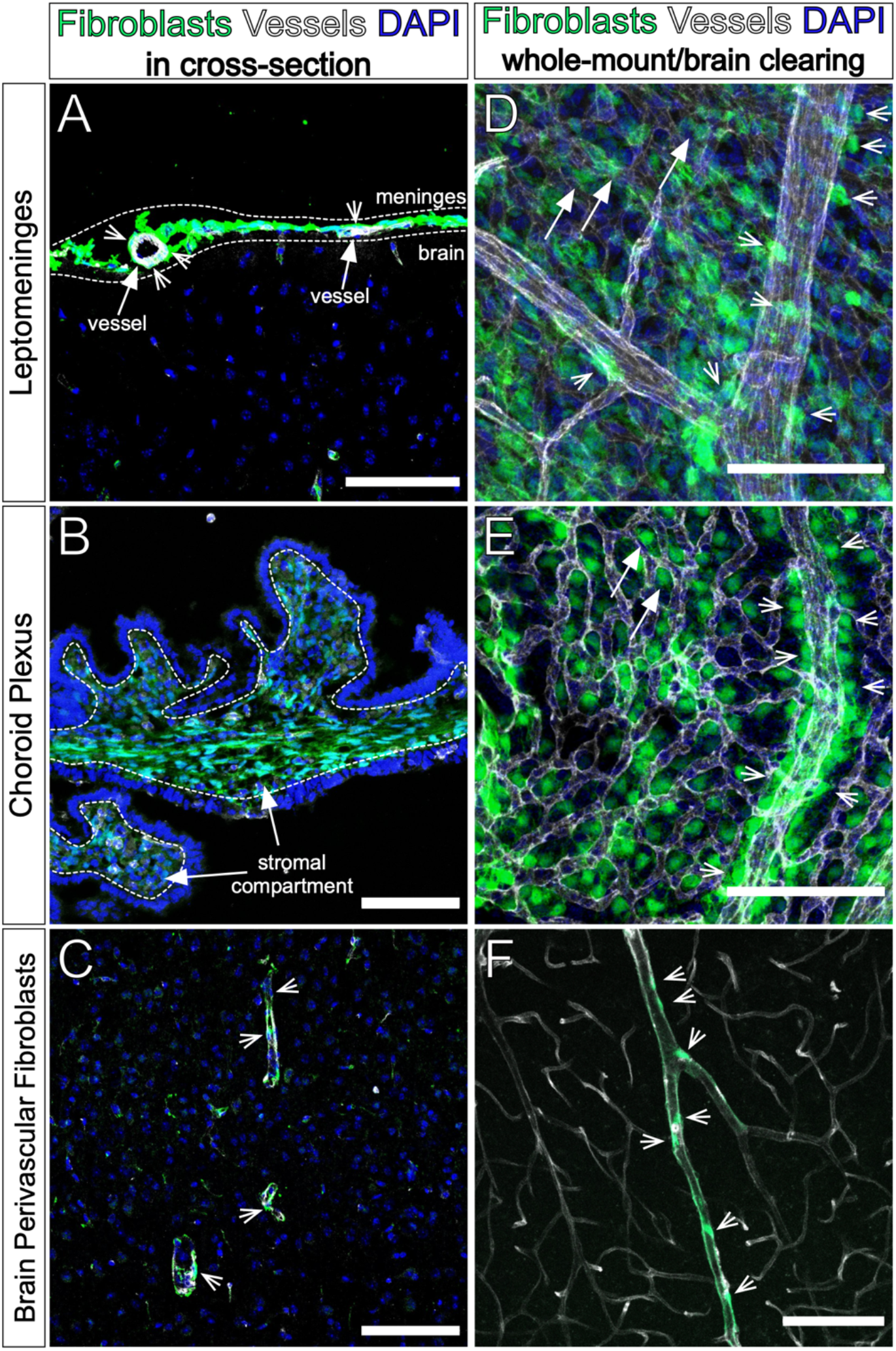
Conventional and improved methods for visualizing CNS fibroblasts. **A)** Adult *Col1a1-GFP* mouse brain section showing cerebral meningeal fibroblasts labeled by GFP (green), vessels labeled by IB4 (white), and DAPI (blue). Meninges shown between dotted lines. Arrows point to vessels (IB4+), arrowheads point to perivascular meningeal fibroblasts. **B)** Postnatal *Col1a1-GFP* mouse brain section showing choroid plexus. Arrows point to stromal compartment, fibroblasts labeled by GFP (green), vessels labeled by IB4 (white), and DAPI (blue). **C)** Adult *Col1a1-GFP* mouse brain section showing fibroblasts labeled by GFP (green) surrounding vessels labeled by IB4 (white), and DAPI (blue). Arrowheads point to perivascular fibroblasts positioned along vessels. **D)** Embryonic day (E)16 *Col1a1-GFP* mouse leptomeninges prepared via whole mount methods, showing fibroblasts labeled by GFP (green), vessels labeled by PECAM (white), and DAPI (blue). Arrowheads point to fibroblasts closely adjacent to a large diameter meningeal vessel. Arrows point to fibroblasts within the vascular plexus. **E)** Postnatal day (P)2 *Col1a1-GFP* mouse lateral ventricle choroid plexus prepared via whole mount methods, showing fibroblasts labeled by GFP (green), vessels labeled by PECAM (white), and DAPI (blue). Arrowheads point to fibroblasts adjacent to a large diameter vessel. Arrows point to fibroblasts interspersed with the vascular plexus. **F)** Adult *Col1a1-GFP* mouse brain cleared with CUBIC protocol, showing fibroblasts labeled by GFP (green) on the outside of vessels labeled with DyLight 649 Tomato Lectin (white). Arrowheads point to perivascular fibroblasts. Scale bars = 100µm.

The meninges are a tri-layered structure (consisting of the pia, arachnoid, and dura layers) that encase the entire CNS, provide protection from physical injury, and contain multiple immune cell types and vascular plexuses that function in neuro-immune surveillance and CSF drainage^9^. Fibroblasts are a major cell type in the meninges and recent single-cell profiling has revealed layer-specific expression profiles of meningeal fibroblasts, potentially related to layer-specific functions^7^. The pia layer, the meningeal layer that forms the basement membrane directly adjacent to the glial limitans of the brain parenchyma, contains an extensive vascular network and a transcriptionally and morphologically distinct fibroblast subtype^7,10,11^. Upon examination of the meninges in a cross-sectional view (Figure 1A), some fibroblasts appear to be perivascular (Figure 1A, arrowheads); this is consistent with prior studies^12,13^. It is unknown if and how these perivascular fibroblasts in the pia layer are a molecularly distinct population from other pial fibroblasts, and from a cross-sectional view it is difficult to appreciate the cellular morphology and the cell-cell interactions occurring between these fibroblasts and the meningeal vasculature (Figure 1A). Improved methods are needed to visualize meninges located perivascular fibroblasts to advance understanding of their potential functions in the meninges.

The choroid plexus is located in each ventricle of the brain and is responsible for the continuous production of cerebrospinal fluid^14,15^. The choroid plexus consists of an outer epithelial layer bound by tight junctions, and an inner stromal compartment consisting of vasculature, immune cells, pericytes/vascular smooth muscle cells, and fibroblasts^15^. While the structure and functions of the choroid plexus epithelial layer is well established^14^, few studies have focused on the diverse cell types that make up the stromal compartment. In particular, there is little information regarding choroid plexus stromal fibroblasts. Examination of the choroid plexus of *Col1a1-GFP* mice in cross-section (Figure 1B) shows fibroblasts in the stromal compartment labeled by GFP. Given their proximity to other cell types in the choroid plexus stromal compartment (vasculature, pericytes/vascular smooth muscle cells, immune cells) and epithelial layer, choroid plexus fibroblasts have the potential to interact with other stromal-located cell types and play key roles in development, homeostasis, as well as disease pathologies. Much like the meninges, the choroid plexus is difficult to examine fully in cross-section (Figure 1B), thus novel methods are required to visualize and appreciate the cell-cell interactions that occur in this structure.

Perivascular fibroblasts reside on the outside of large diameter arterioles and veins within perivascular spaces of the CNS and were first identified for their roles in wound healing and scar formation^3,8,16^. Perivascular fibroblasts are pro-fibrotic in response to acute CNS injuries like stroke, traumatic brain and spinal cord injuries, and undergo transcriptional changes in neuroinflammatory diseases such as multiple sclerosis^3,5,8,16^. Perivascular fibroblasts are also activated in neurodegenerative diseases such as amyotrophic lateral sclerosis, producing factors that damage the blood-brain barrier^4^. How perivascular fibroblasts are activated and contribute to each of these pathologies is an area of ongoing investigation. Additionally, next to nothing is known about the homeostatic role of perivascular fibroblasts within the CNS and how they interact with the vasculature, nor do we know about their developmental origins. Perivascular fibroblasts have conventionally been visualized in cross-sections of the parenchyma (Figure 1C), labeled by GFP in the *Col1a1-GFP* mouse line. Given the complex three-dimensional nature of the brain vasculature, cross-sectional views are not sufficient to appreciate how perivascular fibroblasts interact with vascular and other perivascular cell types in the parenchyma. To answer questions about perivascular fibroblast developmental origins as well as functions in CNS homeostasis, injury, and disease, new visualization techniques are needed.

Here, we detail methods to visualize fibroblasts and blood vasculature located in the meninges, choroid plexus, and perivascular spaces of the CNS. We describe techniques for dissection, immunohistochemistry, whole mounting, and imaging of embryonic, postnatal, and adult meninges and choroid plexus. Whole mount visualization of tissues is not new, however the use of these techniques for examination of these structures is novel. This process results in images that provide a more comprehensive picture of meningeal and choroid plexus fibroblasts and their interactions with the vessels in these structures as well as other resident cell types (Figure 1D-E). We also describe a technique for visualizing perivascular fibroblasts and vasculature in the postnatal and adult brain using optical tissue clearing. Several methods for optical clearing of the CNS have been published^17–20^; we have optimized labeling and imaging the vasculature and perivascular fibroblasts in brains cleared using the CUBIC method. Our protocol results in images that allow for examination of perivascular fibroblast interactions with the brain vasculature in three dimensions (Figure 1F). Together, these methods are useful for revealing details about how fibroblasts in the meninges, choroid plexus, and brain perivascular spaces interact with vessels in these locations. In addition to visualizing fibroblast-vessel interactions, these methods permit descriptions of location and morphology of fibroblasts and other perivascular cell types that is lost in traditional methods. Further, these methods have the potential to be useful for the examination of CNS fibroblast development, functions in homeostasis, and roles in injury and disease.

### 1. Visualizing the embryonic, perinatal, and adult leptomeningeal layers

#### Dissection and fixation of embryonic, postnatal, and adult leptomeninges

Here, we describe methods for dissection of leptomeninges (collective term for combined pia and arachnoid layers) for ages embryonic days (E)14-18 and postnatal days (P)0 through adulthood (Figure 2). All meningeal fibroblasts are labeled by GFP in the *Col1a1-GFP* mouse line. To assay for proliferating cells, we use a EdU assay. EdU is a thymidine analogue that is incorporated into DNA during replication in cells in S-phase, and can be detected using fluorescent probes^21^. If completing EdU detection steps, administer EdU (2.5 mg/mL) via interperitoneial injection prior to anesthetization (embryonic: 150 µL into pregnant dam, P0-P21: 50 µL, adult: 100 µL) (Figure 2A.i-ii.). We allow EdU to circulate for 2 hours, as this is the amount of time it takes for EdU to fully circulate and clear from a mouse’s body. However, note that the following methods are compatible with other assay configurations, including varied circulation time and different cell cycle probes (i.e., BrdU, Ki67). For embryonic timepoints, dissect embryos using established methods from pregnant *Col1a1-GFP* dams and anesthetize using cryoanesthesia until unresponsive, then decapitate (Figure 2A.i.a). For P0-21 and adult mice, euthanize *Col1a1-GFP* mice using age-appropriate methods (Figure 2A.ii.a). To remove blood containing immune cells that can be immunoreactive, mice can be transcardially perfused with PBS (2-5 mL for P0-10, 10mL for P11-21, 20mL for adults) prior to brain dissection. To dissect embryonic brains, make a single incision laterally beginning at the back of the head and move anteriorly towards the eye (Figure 2A.i.b, dotted line). Lift the skin and calvarium dorsally, peeling the tissue away from the surface of the brain (Figure 2A.i.b, arrow). Note that earlier embryos (<E16) will have less ossification and the dura/calvarium will come off as a mesenchymal layer. For postnatal and adult mice, decapitate and cut away the scalp by a midline excision caudal to rostral, from between the ears to immediately superior to the eyes and pull skin laterally to expose the calvarium (Figure 2A.ii.b). Carefully remove the calvarium to maintain the integrity of the dura mater (adhered to the underside of the calvarium) for analyses (if desired) using previously published methods for dura whole mount labeling^22^. To achieve this, insert fine surgical scissors into the foramen magnum and guide them anteriorly around the cortices towards the olfactory bulb (Figure 2A.ii.c). Repeat this process on both sides of the skull, with the excision meeting immediately superior to the olfactory bulb. Lift the calvarium away using forceps by pulling upward from the base of the head, exposing the underlying brain (Figure 2A.ii.c). Using a spatula, gently lift the brain up to remove and place it in a 10 cm plate containing 1X PBS (Figure 2A.i.c and ii.d). Complete the isolation of the leptomeninges using a dissection microscope.

**Figure 2:**
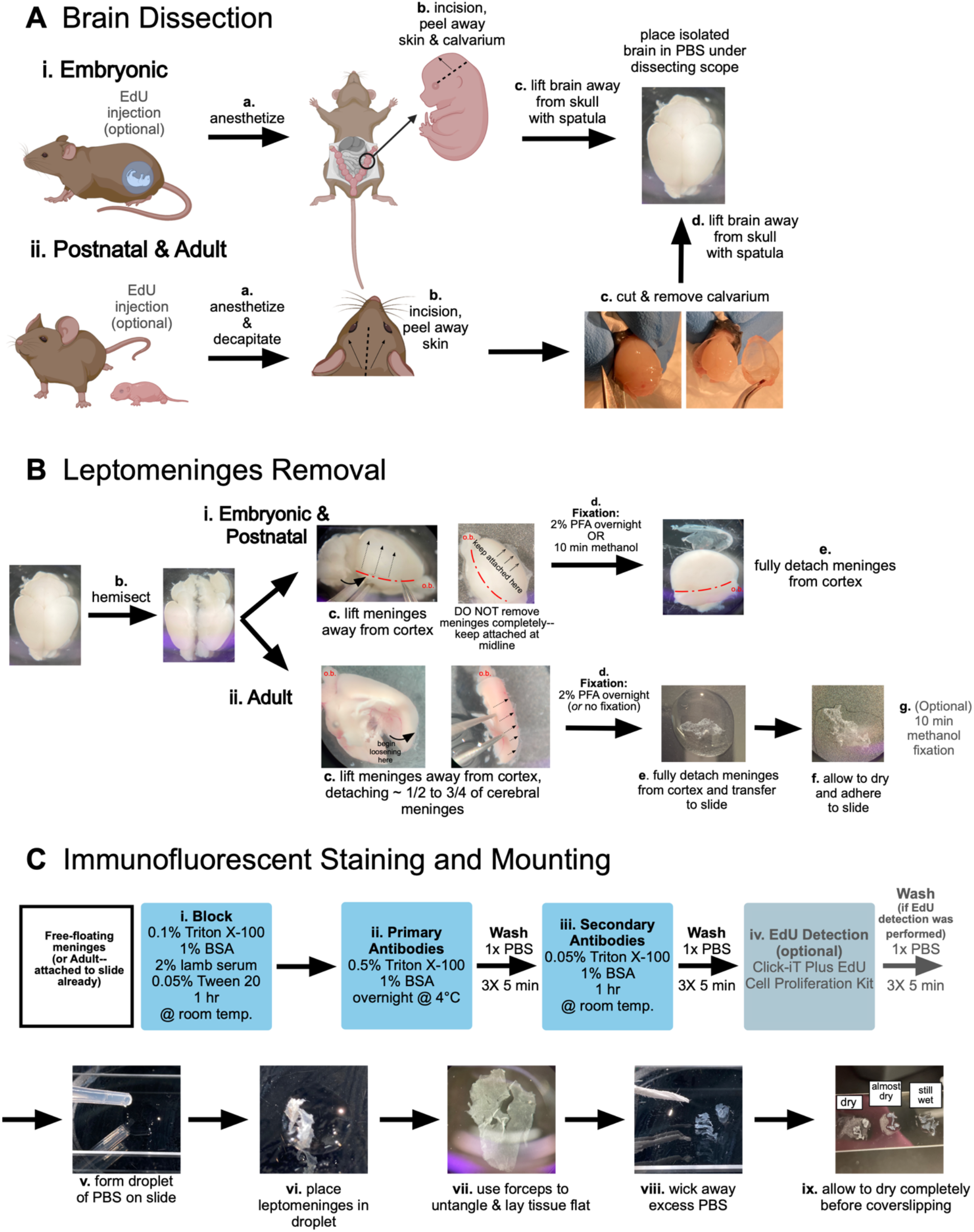
Dissection, fixation, and immunofluorescent staining of embryonic, postnatal, and adult leptomeninges. **A)** Graphical description of brain dissections from (i) embryonic, and (ii) postnatal and adult mice. **B)** Depiction of leptomeninges loosening and removal steps for (i) embryonic and postnatal, and (ii) adult leptomeninges. **C)** Workflow for immunofluorescent staining and mounting of leptomeninges. Brain dissection images are aged postnatal day (P)10 with the exception of 2B.ii.c that is a 9mo old adult.

To loosen the leptomeninges from the surface of the cerebrum, use fine forceps to gently pull the meninges away from the brain (Figure 2B). This is done on both cerebral hemispheres while leaving the meninges attached to the brain near the midline—for fetal and postnatal brain dissections it may help to hemi-sect the brain prior to leptomeninges loosening (Figure 2B.b). Loosening is done to ensure that the leptomeninges do not adhere too tightly to the surface of the brain during the fixation step, permitting removal of the cerebral leptomeninges as an intact structure following fixation. For embryonic and postnatal brains, begin loosening using forceps starting at the lateral incision spot made during the calvarium removal step (Figure 2B.i.c, red dotted line). Note that this incision will not be made on early embryonic brains (<E16) due to the location of the incision. Move the forceps under the meninges and pull them away from the surface of the brain, moving dorsally towards the midline (Figure 2B.i.c, dotted arrows). Do not remove the meninges completely—keep them attached to the brain near the midline (Figure 2B.i.c, arrowheads). Pull the leptomeninges back over the surface of the cerebrum to prevent wrinkles in the leptomeninges. For adult mice, remove the cerebellum and brain stem and hemi-sect the two hemispheres. Remove the thalamus, exposing the cerebrum and hippocampus. Starting at the back of the cerebrum (Figure 2B.ii.c) using fine forceps to slowly loosen a piece of the cerebral leptomeninges that is about ¼ to ¾ of the cerebrum surface. To better visualize the cerebrum surface during meninges removal, cut the hemisphere in the transverse plane and loosen meninges pieces from ventral and dorsal cerebrum separately (Figure 2B.ii.a). Unlike the embryonic and early postnatal brains, the adult leptomeninges will have some hole artifacts made by the loosening process. Of note, this process of leptomeninges removal is especially difficult for brains aged P10 and beyond, possibly due to the leptomeninges becoming thinner with increased brain size, and increasing adherence to the brain. We have found that from ∼P10 to ∼4 months, it is not possible to get large, intact pieces of leptomeninges for whole mount. Instead, it is typical to get small pieces of leptomeninges, usually attached to the middle cerebral artery and primary branches. Once the leptomeninges are loosened, place the brain in 2% paraformaldehyde solution overnight at 4°C then transfer the brain into PBS solution at 4°C (up to 5 days) (Figure 2B.i.d and ii.d).

Methanol fixation is necessary for certain antibodies, in particular junctional proteins found in pial vasculature and arachnoid barrier layer cells (Table 1). Methanol fix can be substituted for 2% PFA with the following modifications. Following leptomeninges ‘loosening’ of embryonic or postnatal brains (described above, Figure 2B), place individual cerebral hemispheres in methanol for 10 min. Brains will turn white. Carefully remove hemispheres with spatula and place in large drop of PBS in the lid of a 10 cm plate. Use fine forceps and dissecting scope, position the hemisphere so that the leptomeninges can be gently detached and floated into the PBS drop. Once detached (Figure 2B.i.e), the leptomeninges are ready for immunostaining (see below, Figure 2C). For adult mice, loosen leptomeninges as described above, detach piece and transfer to a glass slide, then float the leptomeninges into a large drop of PBS so that the ‘brain side’ faces the glass slide (Figure 2B.ii.e). Use a dissection scope and fine forceps to untangle (if necessary) and remove PBS using a transfer pipette and twisted Kimwipe and let dry completely (tissue will appear white when dry). Transfer the slide to plastic container with methanol or use a hydrophobic pen to create a barrier (for fixation on slide); fix for 10min. Transfer slide to PBS and start immunostaining protocol (see below, Figure 2C).

**Table 1:** Markers compatible with meningeal whole mount staining.

| Marker | Type | Cell Type/Region | Ages | Fixation type | Identifier/Reference |
| --- | --- | --- | --- | --- | --- |
| **RALDH2** | Antibody | Arachnoid meningeal fibroblasts, perivascular fibroblasts | All | PFA only | Sigma<br>Cat#: HPA010022 |
| **Collagen1a1-GFP** | Transgenic mouse line | All CNS fibroblasts | All | PFA only | Yata et. al. 2003<br>MGI: 4458034 |
| **NGFR** | Antibody | Pia meningeal fibroblasts | Embryonic, early postnatal | PFA (not tested with methanol) | Cell Signaling Technologies<br>Cat# 8238 |
| **S100a6** | Antibody | Pia meningeal fibroblasts | All | PFA (not tested with methanol) | Novus<br>Cat#: NBP1-89388 |
| **PECAM** | Antibody | Endothelium | All | PFA or Methanol | BD Pharmigen<br>Cat#: 561814 |
| **Desmin** | Antibody | Pericyte/vSMCs | All | PFA (not tested with methanol) | Cell Signaling<br>Cat#: D93F5 |
| **E-cadherin** | Antibody | Arachnoid barrier cells, choroid plexus epithelial cells | All | PFA or Methanol | Becton Dickinson<br>Cat#: 610181 |
| **Claudin 11** | Antibody | Arachnoid barrier cells | All | Methanol only | Invitrogen<br>Cat#: 36-4500 |
| **ZO-1** | Antibody | Endothelium and arachnoid barrier cells | All | Methanol (not tested with PFA) | ThermoFisher<br>Cat#: 33-9100 |
| **CD206** | Antibody | Macrophages | All | PFA (not tested with methanol) | R&D Systems<br>Cat#: AF2535 |
| **Lyve1** | Antibody | Macrophages (leptomeninges & perivascular) | All | PFA (not tested with methanol) | R&D Systems<br>Cat#: AF2125 |
| **VE-Cadherin** | Antibody | Endothelium | All | Methanol only | Abcam<br>Cat#: Ab33168 |
| **Collagen IV** | Antibody | Endothelium, pericytes/vSMC, pia meningeal fibroblasts | All | PFA (not tested with methanol) | BioRad<br>Cat#: 2150-1470 |
| **Iba1** | Antibody | Macrophages | All | PFA (not tested with methanol) | Wako<br>Cat#: 019-19741 |
| **Claudin 5-488** | Antibody | Endothelium | All | PFA or Methanol | ThermoFisher<br>Cat#: 352588 |

Complete leptomeninges isolation of embryonic, postnatal, and adult brains after overnight 2% paraformaldehyde (PFA) fixation by placing brains into a 10 cm dish in 1X PBS solution under a dissection microscope. Detach the leptomeninges from both cerebral hemispheres by gently pulling up on the loosened areas until the meninges can be peeled off as one large piece for staining (Figure 2B i.e). For adult, transfer the piece of leptomeninges to a glass slide containing a large drop of PBS as described above for methanol fixation (Figure 2B.ii.e). Once adult leptomeninges are dried on slide, use hydrophobic pen to create box around meninges piece and proceed with immunostaining protocol.

#### Immunofluorescent staining protocol

We have optimized a protocol for whole mount immunofluorescent staining and EdU detection on embryonic, postnatal, and adult leptomeninges (Figure 2C). Antibodies that have been used successfully with these methods are listed in Table 1. Following dissection of embryonic and postnatal leptomeninges, use fine forceps to transfer each piece into individual wells of a 48 well plate filled with PBS to proceed with staining. For adult leptomeninges, complete staining steps on slides prepared as described above. Recipes for solutions used in this protocol are included in Table 2. Block using a solution consisting of 0.1% Triton X-100, 1% bovine serum albumin (BSA), 2% lamb serum, and 0.05% Tween 20 in PBS on a shaker for one hour at room temperature, adding 100-200µL of blocking solution per well of a 48-well plate (free-floating embryonic or postnatal leptomeninges) or 100-150 µL per piece (on slide, adult leptomeninges) (Figure 2C.i). Following this, incubate free-floating leptomeninges on a shaker overnight at 4°C in 100µL of primary antibodies (see Table 1) diluted with primary solution made up of 1% BSA and 0.5% Triton X-100 in PBS (Figure 2C.ii). Use parafilm to seal plate and prevent evaporation. For adult tissue on slides, perform this step in humidified chamber. After overnight incubation in primary solution, wash the meninges in an excess volume of PBS on a shaker 3 times for 5 minutes each wash. Incubate the meninges on a shaker (embryonic and postnatal) or humidified chamber (adult), protected from light, for 1 hour at room temperature in 100µL of secondary antibodies diluted in secondary solution made up of 1% BSA and 0.05% Triton X-100 (Figure 2C.iii). Repeat PBS washes on a shaker 3 times for 5 minutes per wash. Labeling of cell proliferation via EdU detection is achieved using the Click-iT Plus EdU Alexa Fluor 647 Cell Proliferation Imaging Kit (Thermo Fisher Scientific C10640) (Figure 2C.iv). Following incubation with EdU detection solution (100µL of solution per well of a 48-well plate for free-floating embryonic or postnatal leptomeninges, or 100-150 µL per adult leptomeninges piece mounted on slide), repeat PBS washes on a shaker 3 times for 5 minutes per wash.

**Table 2:** Recipes for reagents used in leptomeningeal and choroid plexus whole mount staining.

| **Solution** | **Components** |
| --- | --- |
| Blocking Solution | 0.1% Triton X-100<br>1% BSA<br>2% Lamb Serum<br>0.05% Tween 20 |
| Primary Solution | 0.5% Triton X-100<br>1% BSA |
| Secondary Solution | 0.05% Triton X-100<br>1% BSA |

After completion of immunostaining and EdU detection, cover-slip adult pieces, or mount embryonic and postnatal leptomeninges on slides following a similar process as described above for adult leptomeninges: transfer leptomeninges to a drop of PBS on a glass slide (Figure 2C.v-vi) and use fine forceps to remove folds and lay the piece of tissue flat with the “brain side” facing the glass (Figure 2C.vii), then use a transfer pipette and Kimwipe to remove excess PBS (Figure 2C.viii). Dry tissue in a covered slide box for approximately 10 to 15 minutes (Figure 2C.ix). Ensure tissue is completely dry (it will appear white) before cover-slipping with a small droplet of mounting media (Fluoromount-G or similar).

Figure 3A depicts images of meninges from embryonic and postnatal *Col1a1-GFP* mice whole mounted using these methods, with vessels labeled for PECAM and EdU detection to label to dividing nuclei (Figure 3A). We are able to clearly distinguish perivascular (Figure 3A, inset i, arrowheads) from non-perivascular (Figure 3A, inset i, arrows) meningeal fibroblasts, and can detect proliferating perivascular cells (Figure 3A, inset i, asterisks). We routinely use these methods with antibody-based staining of meningeal fibroblast markers (full list in Table 1), such as RALDH2 (Figure 3B) and S100a6 (Figure 3C). We also use these methods to label junctional proteins expressed by pial vasculature such as Claudin-5 (Figure 3E) and arachnoid barrier cell enriched junctional proteins E-cadherin (Figure 3E) and Claudin-11 (Figure 3F). Further, leptomeningeal wholemounts can be used to visualize vasculature and meningeal immune cells like border associated macrophages (Cd206+/Lyve1+) (Figure 3G).

**Figure 3:**
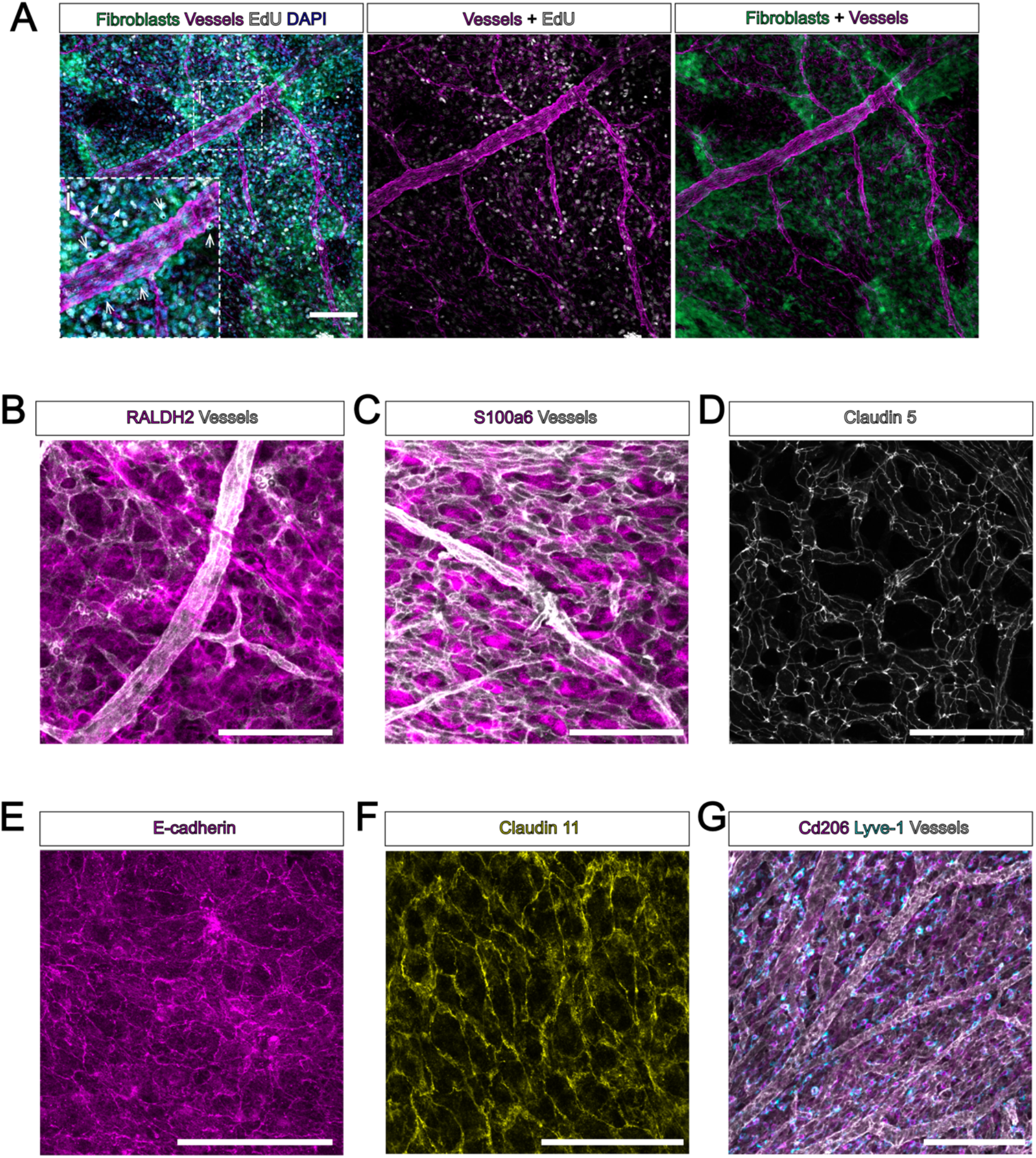
Immunofluorescent staining and EdU labeling of whole mount leptomeninges. **A)** Whole mount leptomeninges from an embryonic day (E)16 *Col1a1-GFP* mouse. Fibroblasts are labeled by GFP (green), vessels labeled by PECAM (magenta), all nuclei labeled by DAPI (Cyan), and proliferating nuclei labeled by EdU (white). Inset **(i)**: arrows indicate non-perivascular fibroblasts labeled by *Col1a1-GFP*, arrowheads indicate perivascular fibroblasts, asterixis next to arrowhead indicates EdU+ cell. **B-C)** Whole mount leptomeninges from embryonic day (E)17 mice. Vessels are labeled by PECAM (white), and meningeal fibroblasts labeled by markers for **B)** arachnoid and perivascular fibroblasts RALDH2 and **C)** pial fibroblasts S100a6 (magenta). **D-F)** Whole mount leptomeninges from postnatal day (P)5-7 mice labeled with junctional markers **D)** endothelial Claudin-5 (white), **E)** arachnoid barrier cell E-cadherin (magenta), and **F)** arachnoid barrier cell Claudin 11 (yellow). **G)** Whole mount leptomeninges from embryonic day (E)17 mice, vessels labeled by PECAM (white) and macrophages labeled by Cd206 (magenta) and Lyve-1 (cyan). Scale bars = 100µm.

### 2. Visualizing the embryonic, postnatal, and adult choroid plexus stroma and vasculature

The brain contains four choroid plexuses, one in each ventricle. Here, we describe methods for dissecting, immunofluorescent staining, and whole mounting of the lateral ventricle choroid plexus for visualization of choroid plexus fibroblasts and vasculature in embryonic, postnatal, and adult mice (Figure 4). This method is compatible with EdU labeling; perform EdU injections as described above. We use the *Col1a1-GFP* mouse line, in which choroid plexus fibroblasts are labeled with GFP. For embryonic choroid plexus isolations (E14-18), euthanize pregnant dams and dissect embryos as for meningeal dissections. For P0-21 and adult mice, euthanize using age-appropriate methods, then mice can be transcardially perfused as described above. Then, remove the brain using methods described for meningeal isolation (Figure 2A). Place the brain in a petri dish containing 1X PBS under a dissecting microscope, use forceps to sagittally hemi-sect the brain and orient it so that the midline is facing upward (Figure 4A-B). From here midline brain structures (hippocampus, thalamus) can be clearly identified, as well as the meninges and choroid plexus. Use one pair of forceps to anchor the thalamus in place and use another other pair in the opposite hand to gently pull the cortex away from the thalamus to reveal the lateral ventricles (Figure 4C, dotted arrows). The choroid plexus will remain attached to brain tissue inside the lateral ventricle (Figure 4C-D, arrows); in perfused tissue it may be difficult to spot, but in un-perfused brains it will have blood in the vasculature and be easier to see. Once the choroid plexus is revealed, use forceps to pinch and tear it away from its attachment at the cortical hem/hippocampus (Figure 4D, asterisk). The midline cerebral meninges and lateral ventricle choroid plexus are continuous structures and may be difficult to distinguish; note that the choroid plexus has a characteristic ‘lettuce leaf” morphology and will appear ‘ruffled’ compared to the meninges (Figure 4D-E). Once the choroid plexus is isolated, use forceps to transfer each piece to individual wells of a 48-well dish containing 100µL of 2% PFA per well and allow to fix as floating pieces for 1 hour at 4°C (Figure 4F). Following fixation, wash three times in 1X PBS for 5 minutes each. From here, proceed with immunohistochemistry, EdU detection and mounting using the same reagents and protocol as for meninges whole mounting (Figure 2C). We show images of choroid plexus from embryonic *Col1a1-GFP* mice whole mounted using the methods described here, with vessels labeled for PECAM (Figure 5A-B) and EdU detection to label proliferating cells (Figure 5B).

**Figure 4:**
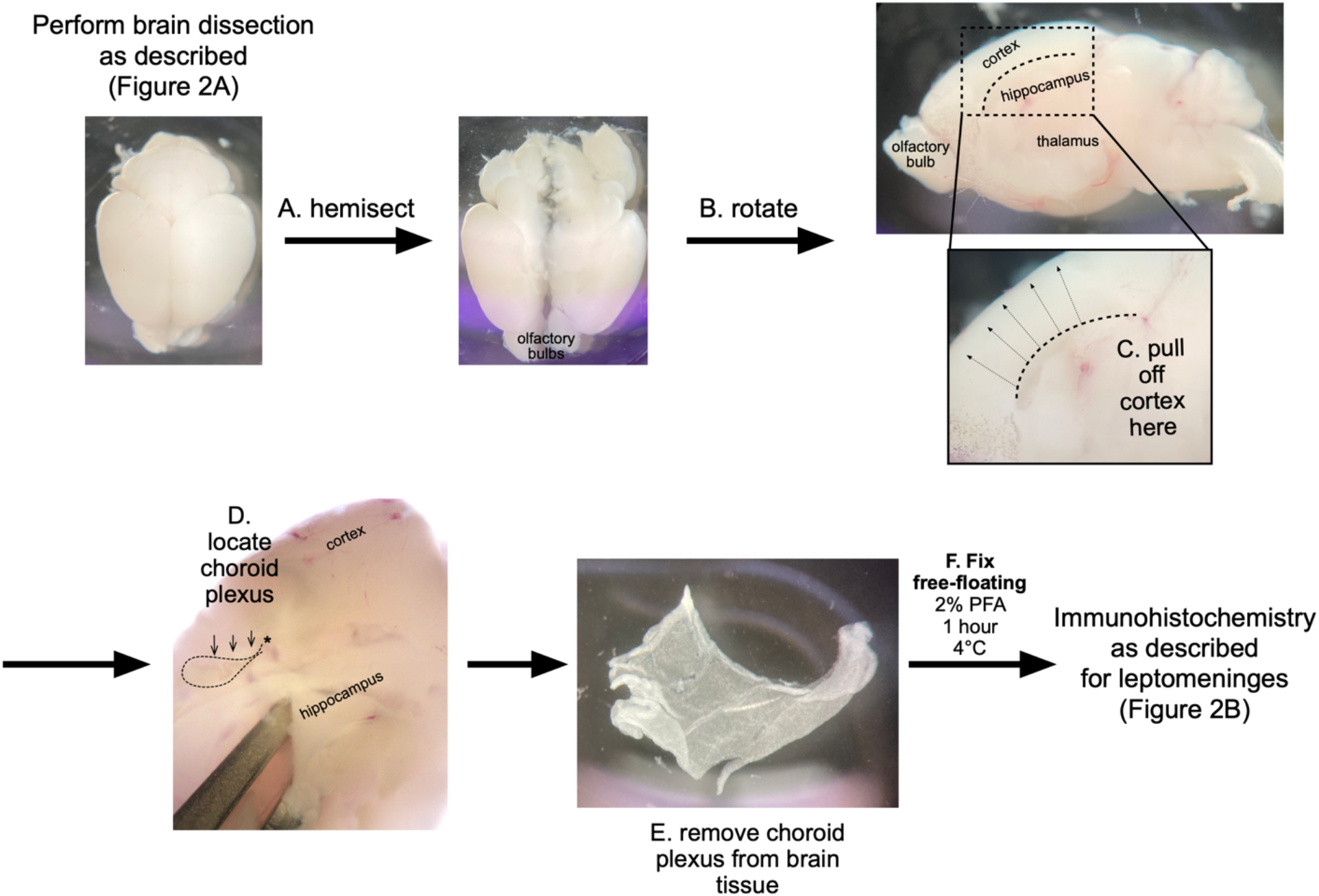
Dissection and whole mounting of lateral ventricle choroid plexus. Workflow for dissection **(A-D)**, removal **(E)**, and fixation **(F)** of lateral ventricle choroid plexus. All brain dissection images are of brains aged postnatal day (P)10 through P17.

**Figure 5:**
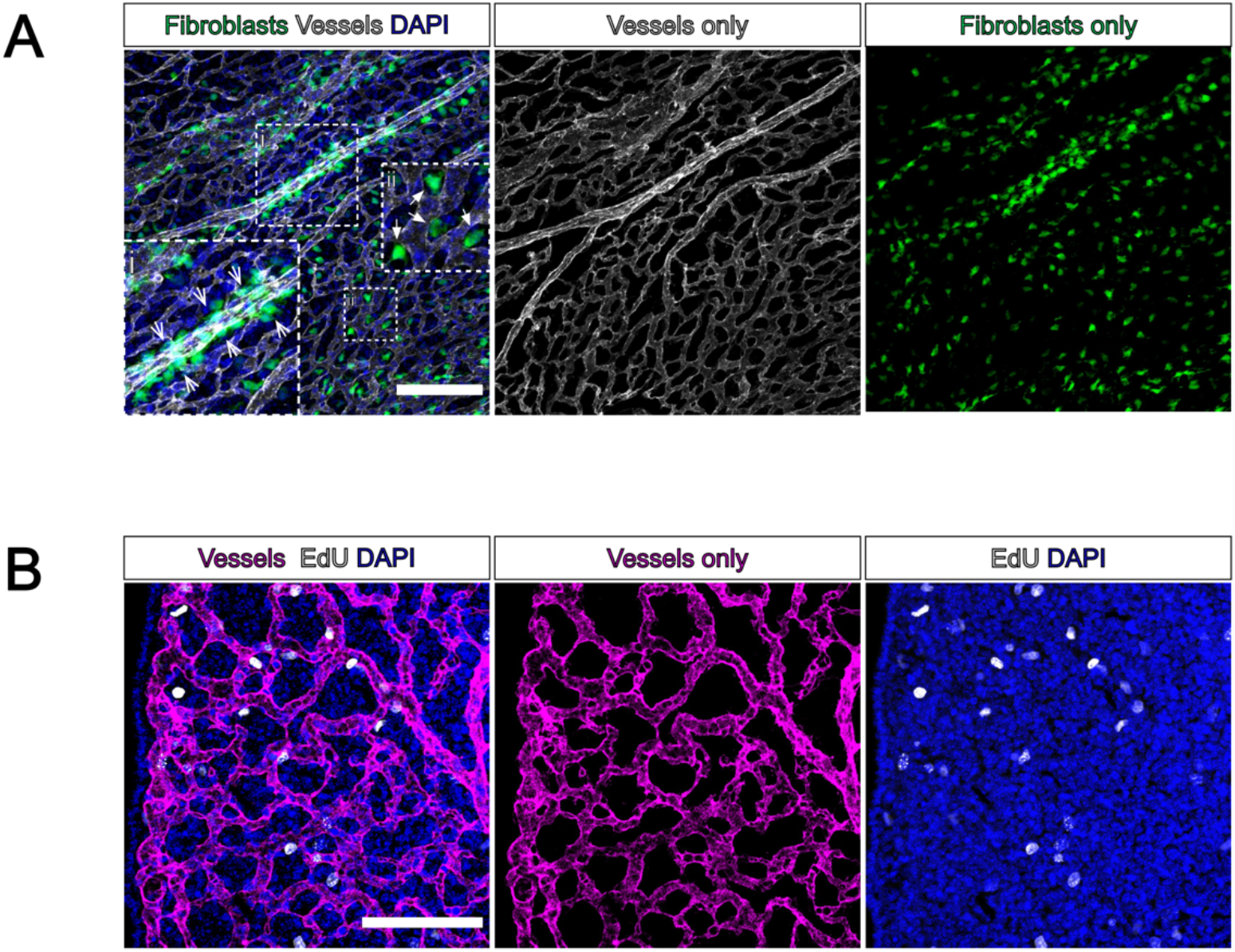
Immunofluorescent staining and EdU labeling of whole mount choroid plexus. Whole mount lateral ventricle choroid plexus from an embryonic day (E)16 *Col1a1-GFP* mouse. **(A)** Fibroblasts labeled by GFP (green), vessels labeled by PECAM (white), and DAPI (blue). Inset (**i)**: Arrowheads indicate perivascular fibroblasts; Inset (ii): arrows indicate non-perivascular fibroblasts. **(B)** EdU is used to visualize proliferating cells in the choroid plexus, nuclei labeled by DAPI (blue), vessels labeled by PECAM (magenta), and proliferating nuclei labeled by EdU (white). Scale bars = 100µm.

### 3. Optical clearing methods for visualization of fibroblast-vessel interactions in the brain

Perivascular fibroblasts are located along large diameter arterioles and venules in the postnatal and adult mouse CNS and are labeled by GFP in the *Col1a1-GFP* mouse line. Here, we describe our optimized protocol for using the CUBIC clearing method for visualizing perivascular fibroblasts in the brain of postnatal and adult *Col1a1-GFP* mice (Figure 6). The CUBIC method is compatible with transgenically-expressed fluorescent proteins as well as fluorescent immunohistochemistry^23,24^, and can be easily adapted. Additionally, we show that this method is compatible with EdU labeling to visualize cell proliferation. First, euthanize and transcardially perfuse mice with 1X PBS then 4% PFA (of each solution: 2-5 mL for P0-10, 10mL for P11-21, 20mL for adults). Dissect brains from mice that were injected with EdU at least 2 hours prior to birth as described above in Figure 2A, paying close attention to keeping all brain regions intact during brain removal. Following brain removal, fix brains overnight in 4% PFA, then wash in 1X PBS at room temperature three times for a duration of 2 hours per wash (6h total) (Figure 6A). Following this step, section brains as desired. For imaging perivascular fibroblasts in the cortex, we section brains embedded in agarose using a vibratome to create 2mm-thick coronal sections (Figure 6B). An alternative approach would be using a precision brain slicer with 1mm slice intervals. Place each brain section into an individual 5mL snap-top tube in 500µL of CUBIC-L solution and incubate at 37°C on a nutator or shaker (Figure 6C). This step and all following steps are done with brains in 5mL snap-top tubes, as this is important to prevent evaporation of solution. Incubate brains in CUBIC-L for at least 2 days, swapping out solution each day for the first 2 days, then every other day until the brains are clear. The brains are sufficiently cleared when they are nearly invisible in solution, or when there appears to be a plateau in the increase of clarity from day to day (Figure 6C, arrow/outline). Note that it may take brains from older mice a longer time to clear, depending on thickness of sections and amount of myelin present. For early postnatal brains, CUBIC-L processing may take as little as 2-4 days, for late postnatal/juvenile mice about 5 days, while for adult mouse brains clearing may take up to 7-10 days.

**Figure 6:**
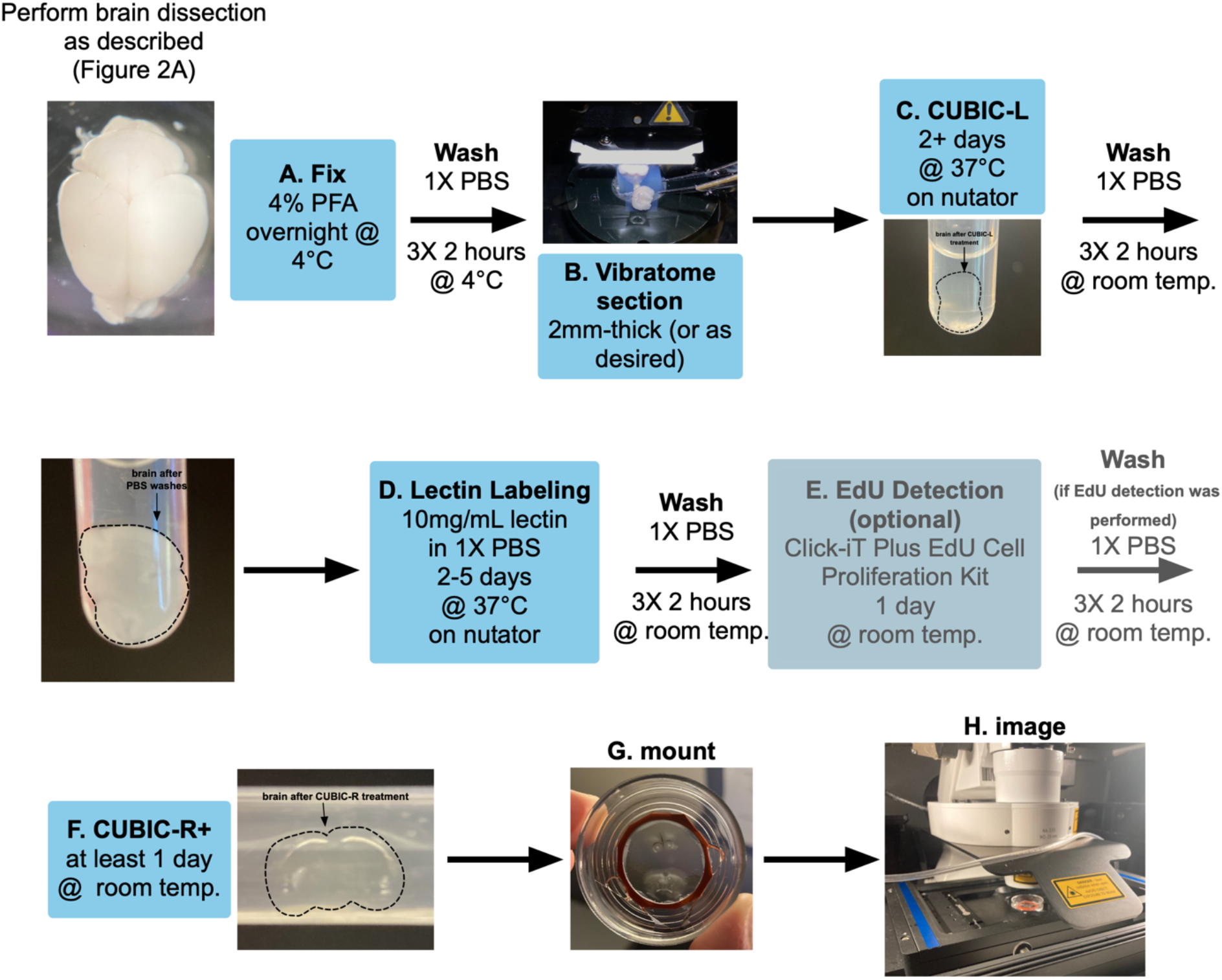
Optimized CUBIC protocol for visualization of brain perivascular fibroblasts. Workflow for preparation **(A-B)**, clearing **(C**,**F)**, lectin vessel labeling **(D)**, and EdU detection (**E)** of *Col1a1-GFP* mouse brains. All brain images are of brains aged postnatal day (P)7 through P14.

Following CUBIC-L incubation, wash brains with an excess volume of 1X PBS at room temperature three times for 2 hours per wash. Following PBS washes, it is typical for sections to appear less clear and cloudy (Figure 6C-D, outline/arrow). Here, we describe steps solely for the labeling of brain vasculature using directly conjugated fluorescent tomato lectin (Vector Laboratories, DL-1178), as well as detection of proliferating cells using EdU. Antibody based immunofluorescent labeling can be performed using published methods^23,24^. Lectin dye is diluted in 1X PBS to a concentration of 10mg/mL, and 500µL of lectin diluted in PBS is added to brains in 5mL snap-top tubes (Figure 6D). Incubate in lectin solution at 37°C on a nutator or shaker for a minimum of 2 days and up to f5 days (Figure 6D). Next, wash with 1X PBS at room temperature three times for 2 hours per wash. If deseired, perform EdU labeling using the Click-iT Plus EdU Alexa Fluor 647 Cell Proliferation Imaging Kit (Figure 6E). Make EdU detection solution as described and incubate brains in snap-top tubes in a minimum of 500µL of solution for 1 day at room temperature while shaking (Figure 6E). Then, wash in 1X PBS at room temperature three times for 2 hours per wash. Finally, place brains into 500µL CUBIC-R+ solution in snap-top tubes at room temperature for at least 1 day (Figure 6F). The brains should revert from an opaque appearance to completely translucent (Figure 6F, arrow/outline).

Brain slices immersed in CUBIC-R+ are imaged in a 35mm glass bottom imaging dish (WPI, FD35-100). Create a 2.5mm-deep well to hold brain slices by trimming and affixing a rubber gasket (Electron Microscopy Sciences, 70336-25) to the bottom of dishes using silicone glue (Figure 6G). For larger slices or whole brains, stack gaskets to create a deeper well. Fill the well created by the gaskets with CUBIC-R+, place and orient brain slices within the well, and carefully place a square coverslip over top, taking care to minimize bubbles. Image brains in dishes as soon as possible using an inverted confocal microscope and objective with a long working distance (Figure 6H). We acquire images on a Zeiss LSM 900 inverted microscope with a 35mm circular stage insert (Figure 6H) using 5x, 10x, and 20x objectives (see Table 3). Here, we show images of large volumes of brain tissue from postnatal *Col1a1-GFP* mice, with vessels labeled using lectin (Figure 7A-B) and dividing nuclei detected via EdU labeling (Figure 7B, arrowheads). Exposure to air will crystallize CUBIC-R+ within a few days—transfer brains to a long-term storage container once imaging is concluded. Brains can be stored in CUBIC-R+ in air-tight containers for up to 6 months while still retaining fluorescent labeling.

**Table 3:** Materials used in leptomeningeal and choroid plexus whole mount staining and CUBIC clearing and imaging of brain slices to visualize perivascular fibroblasts

| Reagent/Resource | Source | Identifier |
| --- | --- | --- |
| Dumont #5SF Forceps | Fine Science Tools | No. 11252-00 |
| Triton X-100 | MP Biomedicals | Cat#: 194854 |
| BSA | VWR | Cat#: 0332 |
| Lamb Serum | ThermoFisher | Cat#: 16070096 |
| Tween 20 | Fisher Bioreagents | Cat#: BP337 |
| PBS | Lonza | Cat#: 17-5170 |
| Click-iT Plus EdU Alexa Fluor 647 Imaging Kit | Thermo Fisher | Ref#: C10640 |
| CUBIC-L | TCI | Cat#: T3740 |
| CUBIC-R | TCI | Cat#: T3741 |
| Tomato Lectin DyLight 649 | Vector Laboratories | Ref#: DL-1178 |
| FluorDish Tissue Culture Dish with Cover Glass Bottom (35mm) | WPI | Cat#: FD35-100 |
| Silicone glue | General Electric | Stock No. SIL2WD |
| Rubber gaskets | Electron Microscopy Sciences | Cat#: 70336-25 |
| Square coverslips | Globe Scientific | Item#: 1404-15 |
| Objective Plan-Apochromat 5x/0.16 M27 | ZEISS | Item#: 420630-9900-000 |
| Objective Plan-Apochromat 10x/0.45 M27 | ZEISS | Item#: 420640-9900-000 |
| Objective LD SC Plan-Apochromat 20x/1.0 Corr M32 85mm | ZEISS | Item#: 421459-9772-000 |
| Click-In System Insert Plate (for 35mm petri dishes) | PeCon GmbH | Item#: 000-440 |

**Figure 7:**
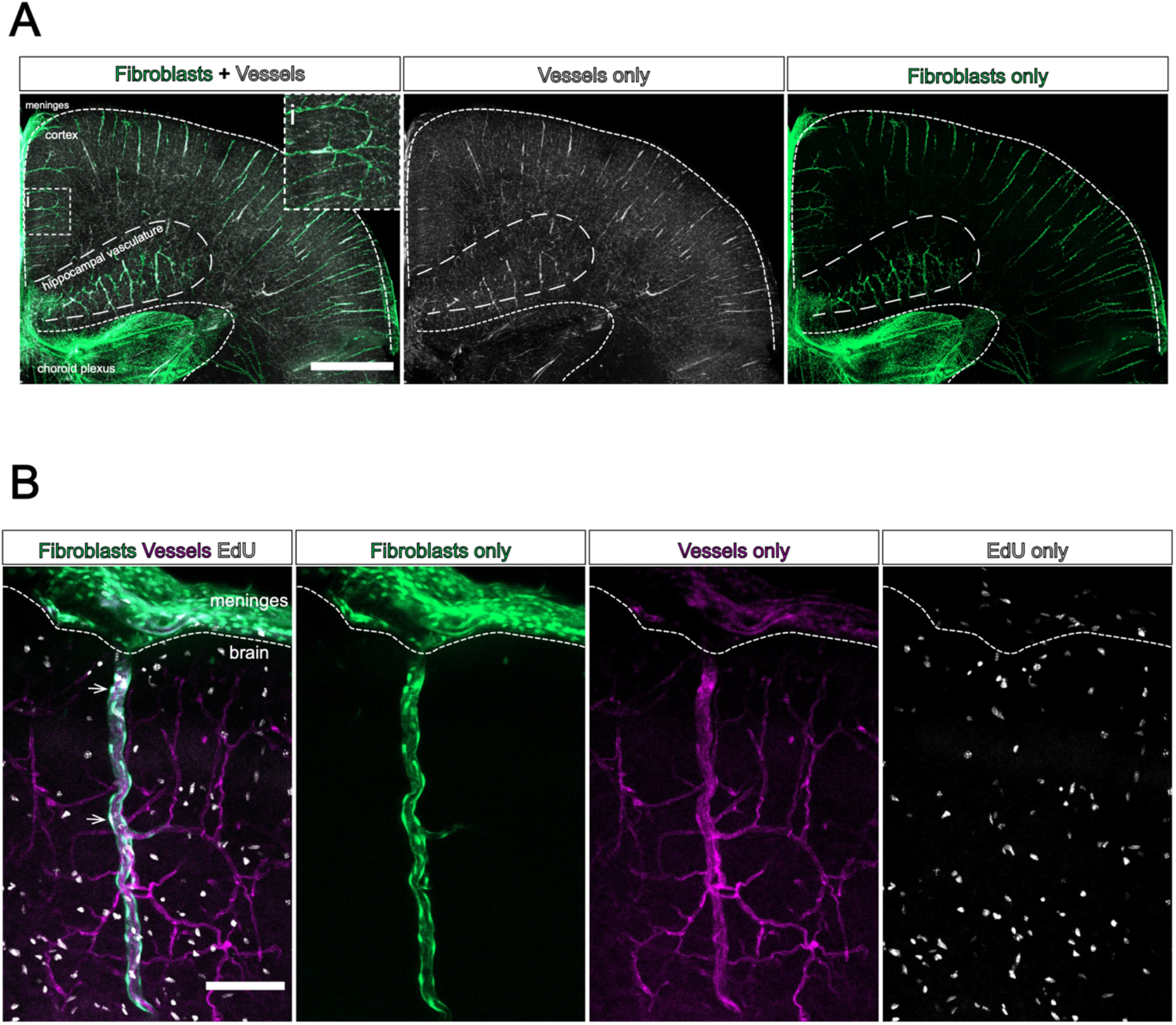
Lectin labeling and EdU detection in CUBIC-cleared *Col1a1-GFP* mouse brains. **(A)** Coronal section of a P10 *Col1a1-GFP* mouse brain, with meninges, cortex, hippocampal vasculature, and lateral ventricle choroid plexus outlined. Fibroblasts are labeled by GFP (green) and vessels are labeled by lectin (white). Maximum projection image created from 5x tile scan, about 300µm depth in the Z-direction. Inset **(i)** shows close-up view of perivascular fibroblasts on a penetrating vessel. Scale bar = 1mm. **(B)** Close-up image of a penetrating vessel in the cortex of a *Col1a1-GFP* mouse brain. Fibroblasts are labeled by GFP (green), vessels are labeled by lectin (magenta), and proliferating cells are labeled by EdU (white). Maximum projection image created from a single-frame 10x scan, about 100µm depth in the Z-direction. Arrowheads indicate proliferating EdU+/GFP+ perivascular fibroblasts. Scale bar = 100µm.

## 4. Discussion

We have developed and optimized methods to image CNS fibroblasts and vasculature in the meninges, choroid plexus, and brain parenchyma that provide substantial benefits over conventional sectioning and labeling methods of these structures. Whole mounting of the meninges and choroid plexus and optical tissue clearing of the brain allows for the examination of the complex vascular plexuses of these two structures and interactions with accompanying fibroblasts that is not possible in thin sectioned tissue. These methods can be readily combined with techniques to label vascular as well as neuronal and glial cell types, making them useful for studying cell-cell interactions of the CNS vascular unit. Further, we show that whole mounting and tissue clearing methods are compatible with EdU labeling and detection making these techniques particularly useful for studying cell cycle dynamics during tissue development, growth, and regeneration.

Recent work has established the molecular identities of meningeal fibroblasts as well as markers specific to each meningeal layer and has suggested distinct functions for the fibroblasts in each layer^7^. The whole mount method we describe here enables us to identify distinct sub-populations of meningeal fibroblasts *in vivo* and to further understand their potential functions and spatial relationship with other meningeal cells, like vascular and immune cell types. Of particular interest are how perivascular fibroblasts in the meninges differ from other meningeal fibroblasts, as it is speculated that perivascular meningeal fibroblasts may be a distinct sub-type of pial fibroblast yet there have been few investigations into perivascular meningeal fibroblast identity. Using whole mount techniques, it will be possible to interrogate how perivascular fibroblasts in the meninges differ from other meningeal cell types in terms of their cellular origins, molecular identity, morphology, and functions. Further, whole mounting will be useful for studying other meningeal cell types that are poorly understood, such as the cells that make up the arachnoid barrier, or rare stem cell populations.

Currently, there are very few insights into the cell types that make up the choroid plexus stromal compartment, consisting of fibroblasts, mural cells, immune cells, and an extensive vascular plexus. A recent single-cell profiling of choroid plexus from each ventricle revealed molecularly distinct subtypes of fibroblasts between the lateral, 3^rd^, and 4^th^ ventricles, suggesting potentially different origins or functions for the fibroblasts in each choroid plexus^25^. However, choroid plexus fibroblasts remain elusive in terms of their cellular origins and their functions. Fibroblasts of the choroid plexus stroma are known to express type-1 collagens and it has been suggested that they developmentally arise from a common lineage as meningeal fibroblasts^26^. There have also been studies describing fibrosis of the choroid plexus in aging, injury, and disease^14,15^, but how choroid plexus fibroblasts contribute to this is unclear. The choroid plexus whole mounting technique described here will be beneficial for investigating the roles of choroid plexus fibroblasts in development, health, aging and disease. It will also be useful for examining how choroid plexus fibroblasts interact with other cell types in the choroid plexus stroma. For example, it has been speculated that choroid plexus fibroblasts may engage with various immune cell types that populate the choroid plexus and thus might play a role in regulating immune infiltration^14,15^; whole mount methods will allow for the close examination of these interactions.

Brain perivascular fibroblasts are a more recently described cell type and have been studied for their roles in injury, neuroinflammation, and neurodegeneration. The developmental origins and homeostatic functions of this cell type are unknown. A prior study from our laboratory established that perivascular fibroblasts are absent from most mouse brain regions at birth, and gradually appear on the outside of blood vessels in the brain through the first three weeks after birth^3^. This work suggested a meningeal origin for perivascular fibroblasts and proposed that perivascular fibroblasts may appear in the brain by migrating along parenchymal blood vessels during postnatal development. Our optimized CUBIC tissue clearing protocol for visualizing perivascular fibroblasts along parenchymal blood vessels is an essential method for investigating this proposed developmental mechanism. Importantly, this method is compatible with fluorescent reporter transgenic mouse lines, making it useful for lineage tracing experiments, and can be integrated with meninges whole mounting methods to fully study the relationship between meningeal and brain perivascular fibroblasts. Further, this method will be useful for visualizing how brain perivascular fibroblasts interact with other parenchymal vascular cell types that reside within the unique perivascular space niche, such as vSMCs, macrophages, and astrocytes, and can be used to study fibroblast-cell interactions during homeostatic conditions as well as in injury and disease.

Owing to advances in sequencing technology, we now know much more about the diversity of fibroblast cell types that reside within the CNS. To advance knowledge of the role of CNS fibroblasts in regulating brain health, it is critical to have context about their locations, size, morphology, interactions with other cell types, and patterns of gene and protein expression. It is likely that CNS fibroblasts participate in many functions and processes that are currently unknown. Whole mounting methods to visualize meningeal and choroid plexus fibroblasts, and brain clearing methods to visualize brain perivascular fibroblasts will allow us to examine these cells in depth and will open new windows for the investigation of their roles in CNS health and disease.

## Acknowledgments

The authors thank Stephanie Bonney for technical work on meninges whole mount protocols. This work is supported by funding from NIH/NINDS (R56 NS098273-06 to JAS and F31 NS125875-01 to HEJ) and NIH/NIGMS 1T32GM141742-01 support for the Developing Scholars summer research program for KAA and HEJ.

